# Glucocorticoid treatment exacerbates mycobacterial infection by reducing the phagocytic capacity of macrophages Glucocorticoids and zebrafish TB

**DOI:** 10.1101/2020.06.19.161653

**Authors:** Yufei Xie, Annemarie H. Meijer, Marcel J.M. Schaaf

## Abstract

Glucocorticoids are effective drugs for treating immune-related diseases, but prolonged therapy is associated with an increased risk of various infectious diseases, including tuberculosis. In this study, we have used a larval zebrafish model for tuberculosis, based on *Mycobacterium marinum* (*Mm*) infection, to study the effect of glucocorticoids. Our results show that the synthetic glucocorticoid beclomethasone increases the bacterial burden and the dissemination of a systemic *Mm* infection. The exacerbated *Mm* infection was associated with a decreased phagocytic activity of macrophages, higher percentages of extracellular bacteria, and a reduced rate of infected cell death, whereas the bactericidal capacity of the macrophages was not affected. The inhibited phagocytic capacity of macrophages was associated with suppression of the transcription of genes involved in phagocytosis in these cells. The decreased bacterial phagocytosis by macrophages was not specific for *Mm*, since it was also observed upon infection with *Salmonella* Typhimurium. In conclusion, our results show that glucocorticoids inhibit the phagocytic activity of macrophages, which may increase the severity of bacterial infections like tuberculosis.

**Summary statement:** Using a zebrafish tuberculosis model, we show that glucocorticoids decrease phagocytosis by macrophages, thereby increasing the bacterial burden. This may explain the glucocorticoid-induced increase in susceptibility to tuberculosis in humans.

## Introduction

Glucocorticoids (GCs) are a class of steroid hormones that are secreted upon stress. The main endogenous GC in our body, cortisol, helps our bodies adapt to stressful situations and for this purpose it regulates a wide variety of systems, like the immune, metabolic, reproductive, cardiovascular and central nervous system. These effects are mediated by an intracellular receptor, the glucocorticoid receptor (GR), which acts as a ligand-activated transcription factor. Synthetic GCs are widely prescribed to treat various immune-related diseases due to their potent suppressive effects on the immune system. However, prolonged therapy with these pleiotropic steroids evokes severe side effects, such as osteoporosis and diabetes mellitus (Buckley and Humphrey, 2018; Suh and Park, 2017). Importantly, the therapeutic immunosuppressive effect of GCs may lead to infectious complications because of the compromised immune system (Caplan et al., 2017; Dixon et al., 2011; Fardet et al., 2016). Similarly, after chronic stress an increased susceptibility to infectious diseases has been observed, due to the high circulating levels of cortisol. In order to better understand these complex effects of GCs, more research is required into how GCs influence the susceptibility to infections and the course of infectious diseases.

Tuberculosis (TB) is the most prevalent bacterial infectious disease in the world, caused by the pathogen *Mycobacterium tuberculosis* (*Mtb*). Despite the efforts made to reach the “End TB Strategy” of the World Health Organization, *Mtb* still infects approximately one-quarter of the world’s population and caused an estimated 1.5 million deaths in 2018, which makes it one of the top 10 causes of death globally (Houben and Dodd, 2016; World Health Organization, 2019). The major characteristic of *Mtb* infection is the formation of granulomas containing infected and non-infected immune cells (Furin et al., 2019). Most *Mtb-*infected people develop a latent, noncontagious infection and do not show any symptoms, with the bacteria remaining inactive, while contained within granulomas (Drain et al., 2018; Lin and Flynn, 2010). About 5-10% of the carriers develop a clinically active TB disease associated with a loss of granuloma integrity (Lin and Flynn, 2010; Parikka et al., 2012). Among those TB patients, the majority manifest a lung infection and around 20% shows infection in other organs like the central nervous system, pleura, urogenital tracts, bones and joints, and lymph nodes (Kulchavenya, 2014). Antibiotics are currently the mainstay for TB treatment, but since antibiotic resistance is rising and an effective vaccine against latent or reactivated TB is still lacking, alternative therapies to control TB are needed (Hawn et al., 2013).

GCs are known to modulate the pathogenesis of TB, but their effects are highly complicated. The use of GCs is considered as a risk factor for TB. Patients who are being treated with GCs have an approximately 5-fold increased risk for developing new TB (Jick et al., 2006), and treatment with a moderate or high dose of GCs is associated with an increased risk of activation of latent TB (Bovornkitti et al., 1960; Kim et al., 1998; Schatz et al., 1976). Consequently, a tuberculin skin test (TST) for screening latent TB is recommended before starting GC therapy (Jick et al., 2006). Moreover, chronic stress which is associated with increased circulating levels of the endogenous GC cortisol, has been shown to be associated with a higher incidence of TB (Lerner, 1996).

Despite the generally detrimental effects of GCs on TB susceptibility and progression, certain types of TB patients are treated with GCs. Chronic TB patients may require GCs for treatment of other disorders, and it has been shown that adjunctive GC therapy may have beneficial effects. Traditionally, adjunctive GC with standard anti-TB therapy has been used for prevention of inflammatory complications in patients with tuberculous meningitis, pericarditis, and pleurisy (Alzeer and FitzGerald, 1993; Evans, 2008; Kadhiravan and Deepanjali, 2010; Singh and Tiwari, 2017). It has been reported that adjunctive GC therapy could improve the probability of survival in tuberculous meningitis and pericarditis (Strang et al., 2004; Thwaites et al., 2004; Torok et al., 2011; Wiysonge et al., 2017). In case of pulmonary TB, the most common form of TB, adjunctive GC therapy is recommended in advanced tuberculosis since broad and significant clinical benefits have been demonstrated (Muthuswamy et al., 1995; Smego and Ahmed, 2003).

Although GCs are being used for adjunctive therapy, the beneficial effects of GC treatment are still under debate. For tuberculous pleurisy TB, the efficacy of GCs is still controversial and for meningitis and pericarditis, information on the GC effects is still incomplete (Prasad et al., 2016; Ryan et al., 2017; Singh and Tiwari, 2017; Wiysonge et al., 2017). A review regarding clinical trials for pulmonary TB showed that, although adjunctive GC therapy appears to have short-term benefits, it is not maintained in the long-term (Critchley et al., 2014). An explanation for the complexity of the effects of GC therapy in TB has been offered by Tobin et al. (2012). They showed that patients suffer from TB as a result of either a failed or an excessive immune response to the mycobacterial infection, and that only the subset of TB meningitis patients with an excessive response, showing a hyperinflammatory phenotype (in their study as a result of a polymorphism in the *LTA4H* gene), benefited from adjunctive GC therapy. It was suggested that GCs may also be beneficial for similar subgroups of patients suffering from other forms of TB (Tobin et al., 2012).

The complex interplay between GC actions and TB underscores the need for a better understanding of the effects of GCs on mycobacterial infection. In the present study we have studied these effects using *Mycobacterium marinum* (*Mm*) infection in zebrafish as a model system. *Mm* is a species closely related to *Mtb* that can infect zebrafish and other cold-blooded animals naturally, causing a TB-like disease (Tobin and Ramakrishnan, 2008). Infection of zebrafish larvae with *Mm* provides an animal model system that mimics hallmark aspects of *Mtb* infection in humans and is widely used for research into mechanisms underlying the course of this disease (Cronan and Tobin, 2014; Meijer, 2016; Ramakrishnan, 2013). Like *Mtb, Mm* is able to survive and replicate within macrophages and, in later stages of infection, induces the formation of granulomas (Davis et al., 2002). The transparency of zebrafish at early life stages makes it possible to perform non-invasive long-term live imaging, which has been used to reveal the earliest stages of granuloma formation (Davis and Ramakrishnan, 2009). In addition, the availability of different transgenic and mutant zebrafish lines and the efficient application of molecular techniques allow us to exploit this zebrafish *Mm* infection model optimally to study both the host factors and bacterial factors involved in mycobacterial infection processes (Meijer, 2016; Tobin and Ramakrishnan, 2008; van Leeuwen et al., 2015). For example, zebrafish studies revealed that infected macrophages can detach from a granuloma and facilitate dissemination to new locations (Davis and Ramakrishnan, 2009). Moreover, the study of an *lta4h* mutant zebrafish line showed that the polymorphism in the *LTA4H* gene is associated with the susceptibility to mycobacterial diseases and the response to adjunctive GC therapy in human, representing a prime example of translational research (Tobin et al., 2012; Tobin et al., 2010).

The zebrafish has proven to be a suitable model for studying the effects of GCs, since the GC signaling pathway is very well conserved between zebrafish and humans. Both humans and zebrafish have a single gene encoding the GR, and the organization of these genes is highly similar (Alsop and Vijayan, 2008; Schaaf et al., 2009; Stolte et al., 2006). Both the human and the zebrafish gene encodes two splice variants, the α-isoform, the canonical receptor, and the β-isoform, which has no transcriptional activity (Schaaf et al., 2009). The DNA binding domain (DBD) and ligand binding domain (LBD) of the canonical α-isoform of the human and zebrafish GR share similarities of 98.4% and 86.5% respectively (Schaaf et al., 2009). The zebrafish GR α-isoform, hereafter referred to as Gr, mediates GC effects that have traditionally been observed in humans and other mammals as well, like the effects on metabolism (Chatzopoulou et al., 2015) and the suppression of the immune system (Chatzopoulou et al., 2016). This makes the zebrafish an ideal model to study the mechanisms of GC action *in vivo* (Facchinello et al., 2017; Faught and Vijayan, 2019). In a recent study, we have demonstrated that GC treatment inhibits the activation of the immune system in zebrafish larvae upon wounding (Xie et al., 2019). The migration of the neutrophils and the differentiation of macrophages was attenuated upon treatment with the synthetic GC.

In the present study, to investigate the functional consequences of the previously observed GC effects on immune cells, we have investigated how GCs modulate the course of an *Mm* infection in zebrafish larvae. We demonstrate that beclomethasone increases the level of *Mm* infection and tissue dissemination. This increased *Mm* infection can be explained by an inhibition of the phagocytic activity of macrophages by beclomethasone, which did not affect the microbicidal capacity of these cells. The inhibitory effect of beclomethasone on phagocytosis, which most likely results from Gr interfering with the transcription of genes required for phagocytosis, results in a higher percentage of extracellular bacteria, which eventually leads to an exacerbation of the *Mm* infection.

## Results

### Beclomethasone increases mycobacterial infection through Glucocorticoid receptor (Gr) activation

To study the effect of GC treatment on *Mm* infection in zebrafish, we pretreated zebrafish embryos with beclomethasone and infected them intravenously with fluorescently labelled *Mm*. At 4 days post infection (dpi), the bacterial burden was assessed by quantification of pixel intensities of fluorescence microscopy images. We found that the bacterial burden increased by 2.3 fold when embryos were treated with 25 <u>μM</u> beclomethasone compared with the vehicle-treated group (Figure 1 A, C). Beclomethasone treatment at lower concentrations of 0.04, 0.2, 1 and 5 <u>μM</u> did not affect the bacterial burden. Therefore, a concentration of 25 μM beclomethasone was used in subsequent experiments. We have previously shown that this concentration effectively reduces wound-induced leukocyte migration in zebrafish as well (Xie et al., 2019).

**Figure 1.**
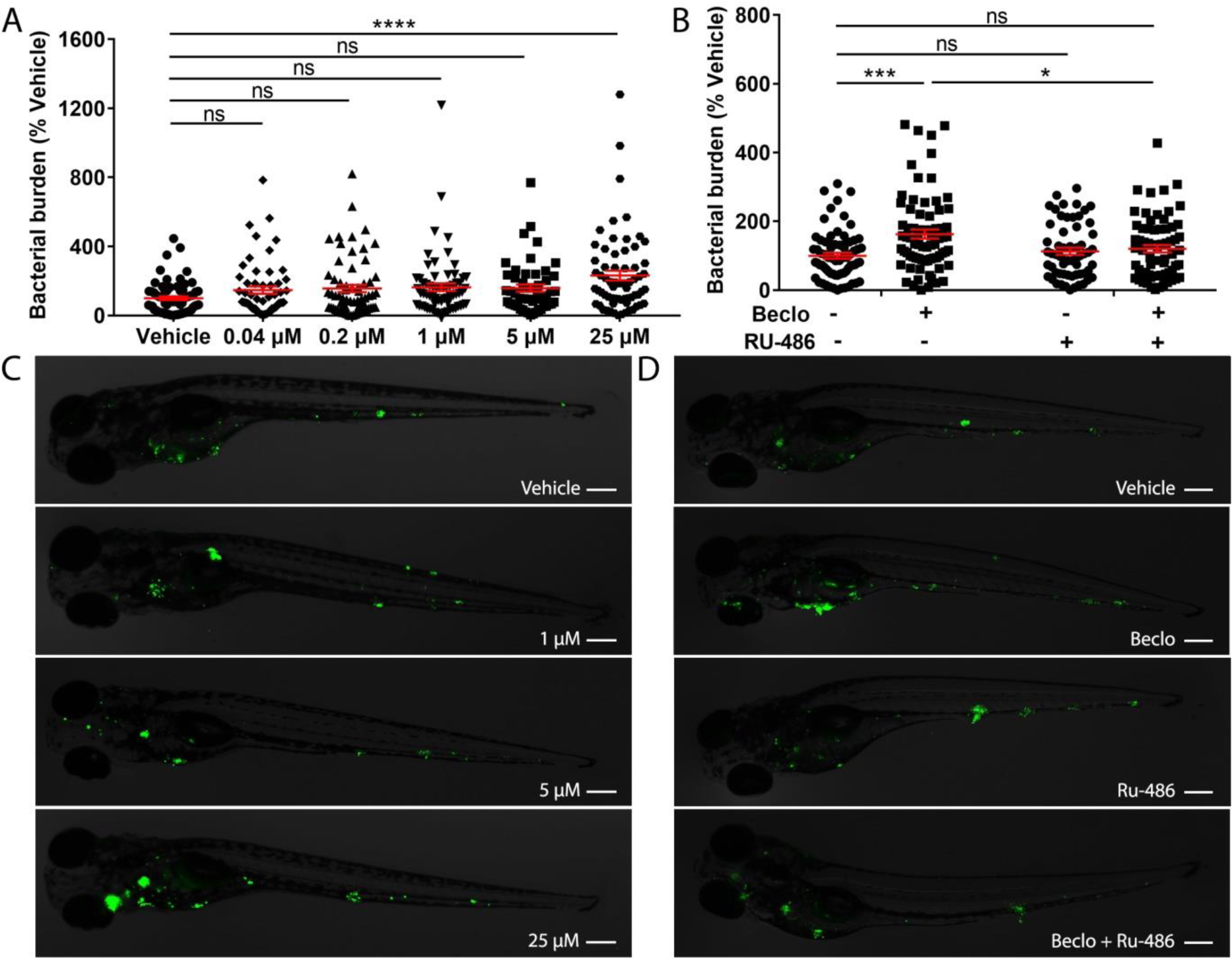
Effect of beclomethasone on *Mm* infection burden in zebrafish. A. Bacterial burden of zebrafish larvae at four days after intravenous injection (at 28 hpf) of *Mm* and treatment with vehicle or different concentrations of beclomethasone (beclo), started at 2 h before infection. Statistical analysis by one-way ANOVA with Bonferroni’s post hoc test revealed that the bacterial burden was significantly increased in the group treated with 25 μM beclomethasone, compared to the burden of the vehicle-treated group. B. Effect of the GR antagonist RU-486 on the beclomethasone-induced increase of the bacterial burden at 4 dpi. The bacterial burden was significantly increased by beclomethasone (25 μM) treatment and this increase was abolished in the presence of RU-486. Statistical analysis was performed by two-way ANOVA with Tukey’s post hoc test. In panels A and B, each data point represents a single larva and the means ± s.e.m. of data accumulated from three independent experiments are shown in red. Statistical significance is indicated by: ns, non-significant; * P<0.05; *** P<0.001; **** P<0.0001. C-D. Representative fluorescence microscopy images of *Mm*-infected larvae at 4 days post infection (dpi), representing experimental groups presented in panels A and B. Bacteria are shown in green. Scale bar = 200 μm.

To demonstrate that the beclomethasone-induced increase in bacterial burden was not due to a general toxicity of beclomethasone but mediated specifically by the Gr, we used the GR antagonist RU-486. The results of these experiments showed that the beclomethasone-induced increase in bacterial burden at 4 dpi was abolished when co-treatment with RU-486 was applied (Figure 1 B, D), which indicates that the effect of beclomethasone requires activation of Gr. No significant difference was observed when the RU-486-treated larvae were compared to the vehicle-treated group. In conclusion, beclomethasone increases the level of *Mm* infection in zebrafish larvae and this effect is mediated by Gr.

### Beclomethasone treatment leads to a higher infection and dissemination level without influencing the microbicidal capacity of macrophages

Subsequently, we analyzed the effect of beclomethasone on *Mm* infection in more detail. The total bacterial burden (Figure 2 A), the number of bacterial clusters per individual (Figure 2 B) and the average size of the bacterial clusters (Figure 2 C) were quantified at 1, 2, 3 and 4 dpi. The results showed that the difference in bacterial burden between the beclomethasone-treated group and the vehicle group was not significant at 1-3 dpi, but that a significant difference was observed at 4 dpi (6186.1±626.5 vs 2870.5±235.0). However, a significant increase in the number of bacterial clusters in the beclomethasone-treated group was already detected at 3 dpi (28.3±1.9 vs 18.1±1.5 in the vehicle group) which was sustained at 4 dpi (64.2±3.5 vs 35.4±2.6). The size of the bacterial clusters at 4 dpi was also increased in the beclomethasone-treated group compared to the cluster size in the vehicle-treated group (741.6±58.3 vs 498.3±45.7). The increase in the number of bacterial clusters indicates an increased dissemination of the infection due to beclomethasone treatment. We confirmed this effect of beclomethasone on bacterial dissemination using hindbrain infection (Figure 2 D, E). Following *Mm* injection into the hindbrain ventricle, 66.1±2.0% of embryos in the vehicle-treated group showed disseminated infection in tissues of the head and tail at 24 hours post infection (hpi), while a significantly higher number (76.4±2.6%) showed this dissemination in the beclomethasone-treated group.

**Figure 2.**
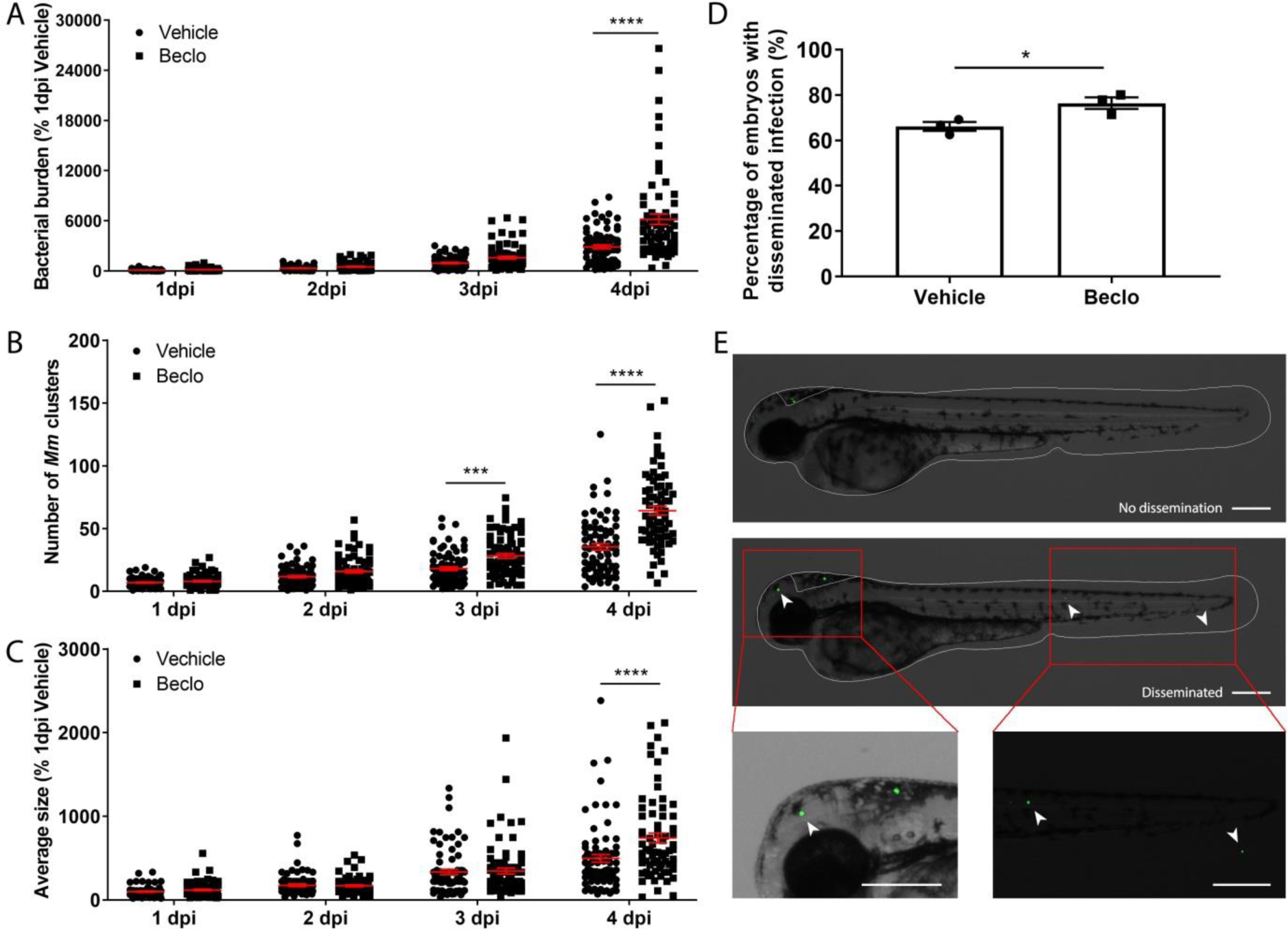
Beclomethasone effects on *Mm* infection progression and bacterial dissemination. A-C. Bacterial burden (A), number of bacterial clusters (B) and the average size of bacterial clusters (C) were determined at 1, 2, 3 and 4 dpi following intravenous *Mm* injection (28 hpf) and treatment with vehicle or 25 μM beclomethasone, started at 2 h before infection. Significant increases due to the beclomethasone treatment were observed for all parameters at 4 dpi. For the number of bacterial clusters, the increase was also significant at 3 dpi. Statistical analysis was performed by two-way ANOVA with Tukey’s post hoc test. Each data point represents a single larva and the means ± s.e.m. of data accumulated from three independent experiments are shown in red. Statistical significance is indicated by: *** P<0.001; **** P<0.0001. D. Effect of beclomethasone on dissemination of *Mm* by hindbrain ventricle injection. Hindbrain infections were performed at 28 hpf, and at 24 hours post infection (hpi), a significantly increased percentage of larvae with disseminated Mm infection was detected in the beclomethasone-treated group compared to the vehicle group. Statistical analysis was performed by two-tailed t-test. Values shown are the means ± s.e.m. of three independent experiments with a total sample size of 27 in the vehicle-treated group and 31 in the beclomethasone-treated group. Statistical significance is indicated by: * P<0.05. E. Representative images of embryos with and without dissemination of the infection upon hindbrain injection of *Mm*. Scale bar = 200 μm.

To study whether the increased infection and dissemination was related to the microbicidal capacity of macrophages, we injected *Mm Δerp* bacteria which are deficient for growth inside macrophages (Clay et al., 2008). No significant difference was observed for the number of *Mm* clusters (Figure 3A) and the percentage of *Mm* inside macrophage (Figure 3B) between the beclomethasone-treated group and the vehicle-treated group. To assess the ability of macrophages to kill bacteria, we quantitated the percentage of bacteria-containing macrophages that contained only 1-10 bacteria in the tail region at 44 hpi (Sommer et al., 2020). There was no significant difference in this percentage between the vehicle-treated group (82.0±4.9%) and the beclomethasone-treated group (81.6±5.0%) (Figure 3C-E). Taken together, these findings indicate that beclomethasone treatment leads to a higher overall *Mm* infection level and increased dissemination, and that these effects are not related to an altered microbicidal capacity of macrophages.

**Figure 3.**
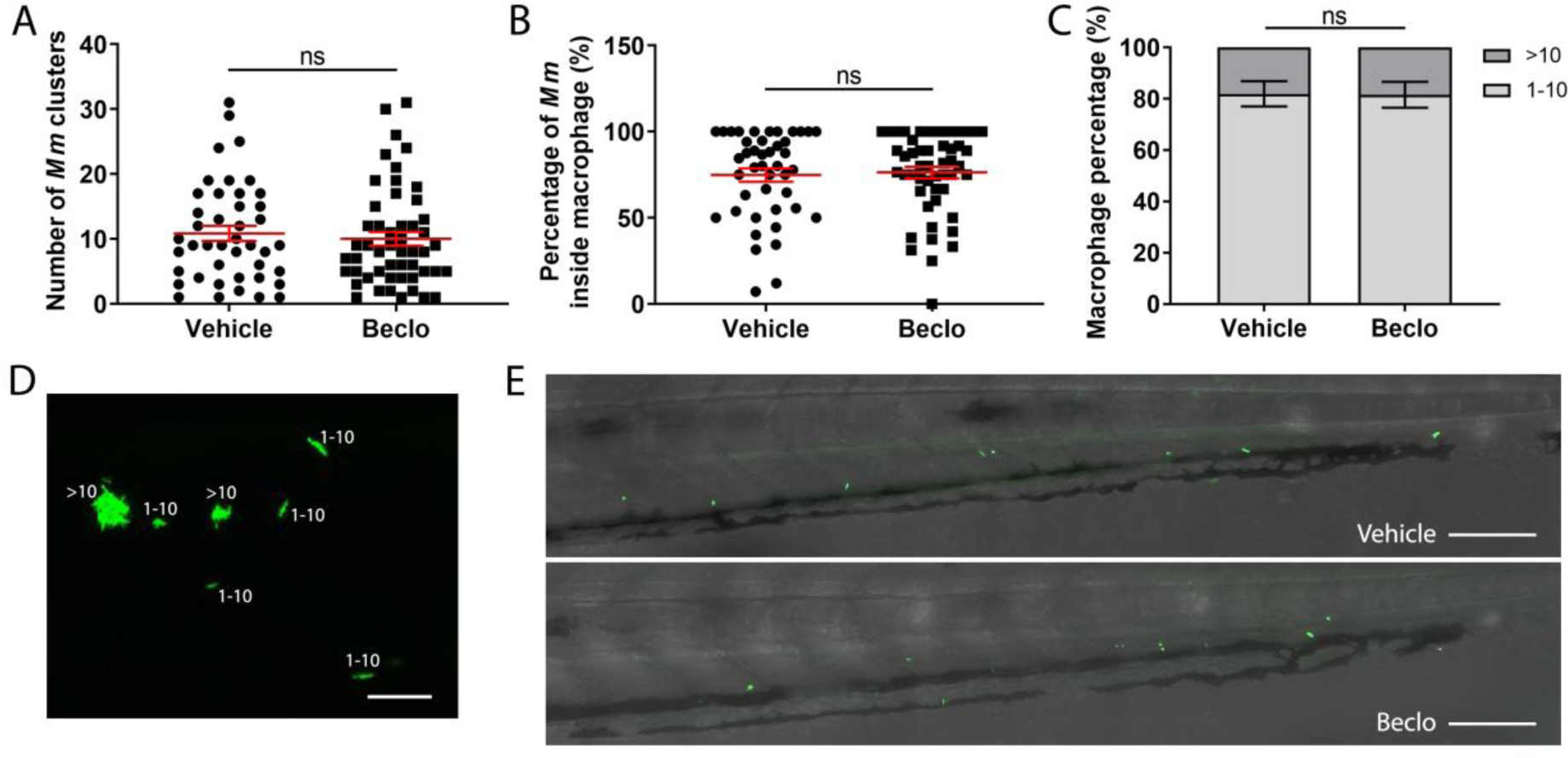
Effect of beclomethasone on *Mm Δerp* mutant bacterial growth. A-C. The *Mm Δerp* mutant strain was injected intravenously at 28 hpf, and at 44 hpi the number of *Mm* clusters (A) and the percentage of *Mm* inside macrophages (B), and the percentage of macrophages that contained 1-10 or more than 10 bacteria (of all macrophages containing bacteria) (C) were determined. No significant difference was observed between the vehicle- and beclomethasone-treated groups. Statistical analysis was performed using two-tailed t-tests. Values shown are the means ± s.e.m. of three independent experiments, with each data point representing a single embryo. Statistical significance is indicated by: ns, non-significant. D. Representative confocal microscopy image of *Mm Δerp* bacterial clusters (bacteria in green), indicated are clusters containing 1-10 bacteria and clusters containing more than 10 bacteria. Scale bar = 20 μm. E. Representative images of the tail regions of a vehicle- and a beclomethasone-treated embryo infected with *Mm Δerp* bacteria. Scale bar = 100 μm.

### Beclomethasone activation of Gr inhibits macrophage phagocytic activity

Since previous studies showed that increased *Mm* infection could be related to decreased phagocytic activity of macrophages in zebrafish (Benard et al., 2014), we studied the effect of beclomethasone on phagocytosis. We used the *Tg(mpeg1:mCherry-F)* line in which macrophages are fluorescently labeled, and assessed phagocytic activity of macrophages by determining the percentage of *Mm* that were internalized by macrophages in the yolk sac area (Benard et al., 2014) (Figure 4 A-C). In the vehicle-treated group, the percentage of phagocytosed *Mm* was 17.4±3.5% at 5 minutes post infection (mpi) and gradually increased to 41.9±4.9% and 52.8±5.2% at 15 and 25 mpi respectively. At each of these time points, a lower percentage of *Mm* were phagocytosed in the beclomethasone-treated group (4.6±1.6% at 5 mpi, 25.7±4.7% at 15 mpi and 34.0±5.2% at 25 mpi). In addition, we studied the involvement of Gr in the beclomethasone-induced inhibition of phagocytosis at 5 mpi, by co-treatment with the GR antagonist RU-486 (Figure 4 D). We found that the decreased phagocytic activity that was observed upon beclomethasone treatment was abolished when larvae were co-treated with RU-486, indicating that the inhibition of phagocytosis by beclomethasone is mediated by Gr.

**Figure 4.**
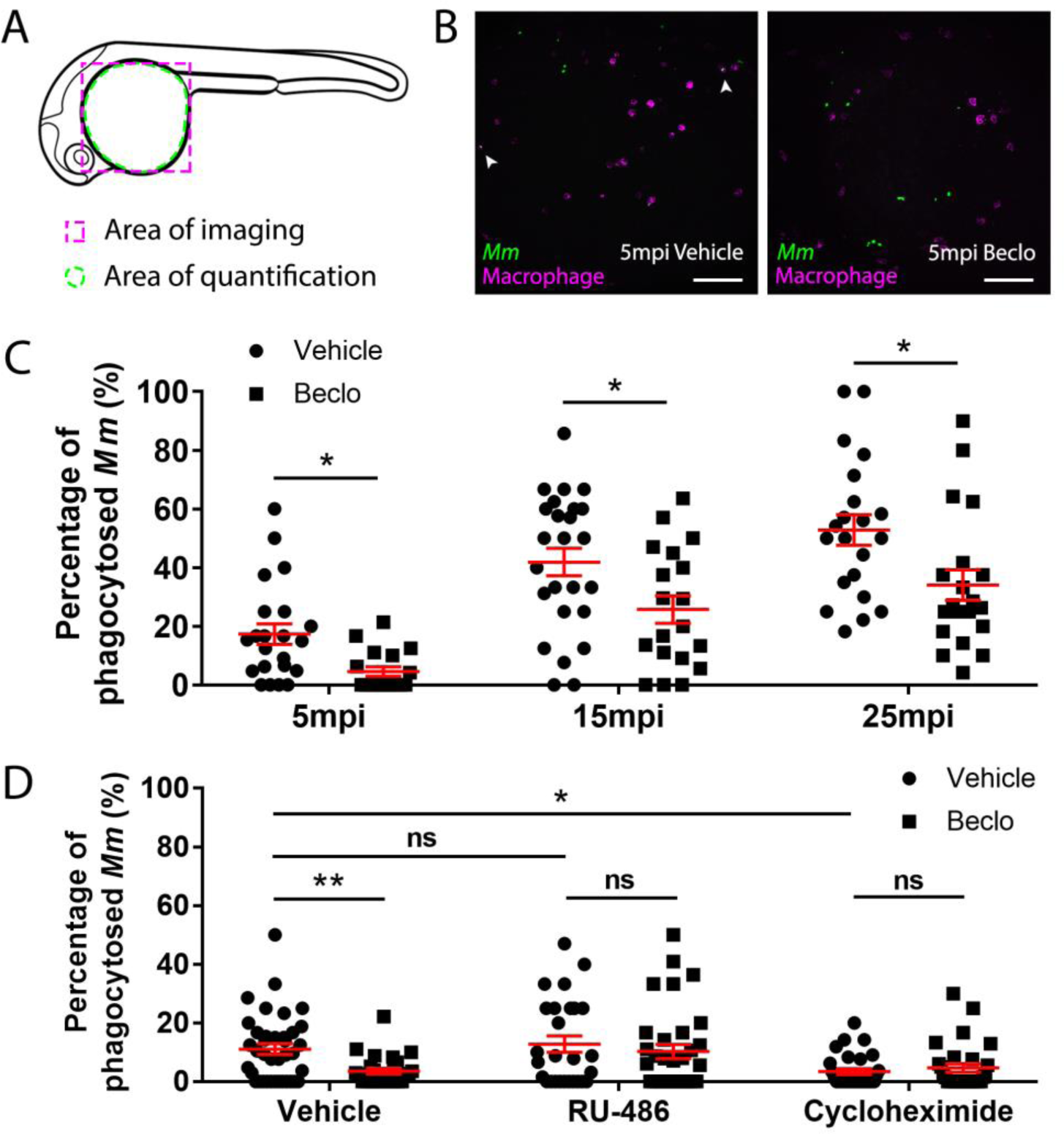
Effect of beclomethasone on phagocytic activity of macrophages and its dependency on Gr and *de novo* protein synthesis. A. Schematic drawing of a zebrafish embryos at 28 hpf indicating the areas of imaging (purple dashed box, used for representative images) and quantification (green dashed circle) of *Mm* phagocytosis. B. Representative confocal microscopy images of embryos of the *Tg(mpeg1:GFP)* line injected with Mm at 28 hpf. Images were taken of infected embryos that were vehicle- or beclomethasone-treated at 5 minutes post infection (mpi). Macrophages are shown in magenta, bacteria in green. Scale bar = 100 μm. Arrowheads indicate bacterial clusters phagocytosed by macrophages. C. Percentages of phagocytosed *Mm* clusters (of total number of *Mm* clusters) at 5, 15 and 25 mpi. Statistical analysis, performed by fitting data to a beta inflated regression with Tukey’s post hoc test, showed that beclomethasone decreased this percentage at all three time points. D. Effects of RU-486 and cycloheximide on the beclomethasone-inhibited phagocytic activity. Embryos were treated with vehicle or beclomethasone and received either a vehicle, RU-486 or cycloheximide co-treatment two hours before injection of *Mm* at 28 hpf, and phagocytic activity was determined at 5 mpi. The significant inhibitory effect of beclomethasone on phagocytosis was not observed in the presence of RU-486. Cycloheximide, just like beclomethasone, significantly inhibited the phagocytic activity, and the combined cycloheximide /beclomethasone treatment showed the same level of inhibition. Statistical analysis was performed by fitting data to a beta inflated regression with Tukey’s post hoc test. In panels C and D, each data point represents a single embryo and the means ± s.e.m. of data accumulated from three independent experiments are shown in red. Statistical significance is indicated by: ns, non-significant; * P<0.05; ** P<0.01.

Gr generally acts as a transcription factor, modulating the transcription rate of a wide variety of genes. To study whether phagocytosis could be modulated by altering the process of protein synthesis, we blocked *de novo* protein synthesis by treatment with cycloheximide (Figure 4 D). We observed that the phagocytic activity of macrophages at 5 mpi was decreased by the cycloheximide treatment (3.4±1.0% vs 11.1±1.8% in the vehicle group). These data demonstrate that phagocytosis depends on *de novo* protein synthesis, and suggest that modulating transcription could be the mechanism underlying the inhibition of phagocytosis by Gr.

### Beclomethasone treatment results in fewer intracellular bacteria and limits infected cell death

To further analyze the possible mechanisms underlying the beclomethasone-induced increase in the *Mm* infection level, we assessed the percentage of bacteria that are present inside and outside macrophages in the caudal hematopoietic tissue (CHT) at 48 hpi using *Mm* infection in the *Tg(mpeg1:GFP)* line. The results showed that beclomethasone treatment resulted in a decreased percentage of intracellular bacteria (23.8±3.0%) compared to the percentage in the vehicle-treated group (36.5±3.6%) (Figure 5 A, C). This result was in line with the observed decrease in phagocytosis at earlier stages of infection. Finally, we used terminal deoxynucleotidyl transferase dUTP nick end labelling (TUNEL) staining to detect cell death, and we performed this staining at 48 hpi (Zhang et al., 2019). In the beclomethasone-treated group, the percentage of *Mm* that were colocalized with TUNEL staining (9.4±1.6%) was significantly lower compared to the percentage of the vehicle group (17.2±2.3%) (Figure 5 B, D). These data suggest that the observed inhibition of phagocytosis upon beclomethasone treatment causes a decrease in the percentage of intracellular bacteria, which underlies the lower numbers of macrophages undergoing cell death as a result of the *Mm* infection.

**Figure 5.**
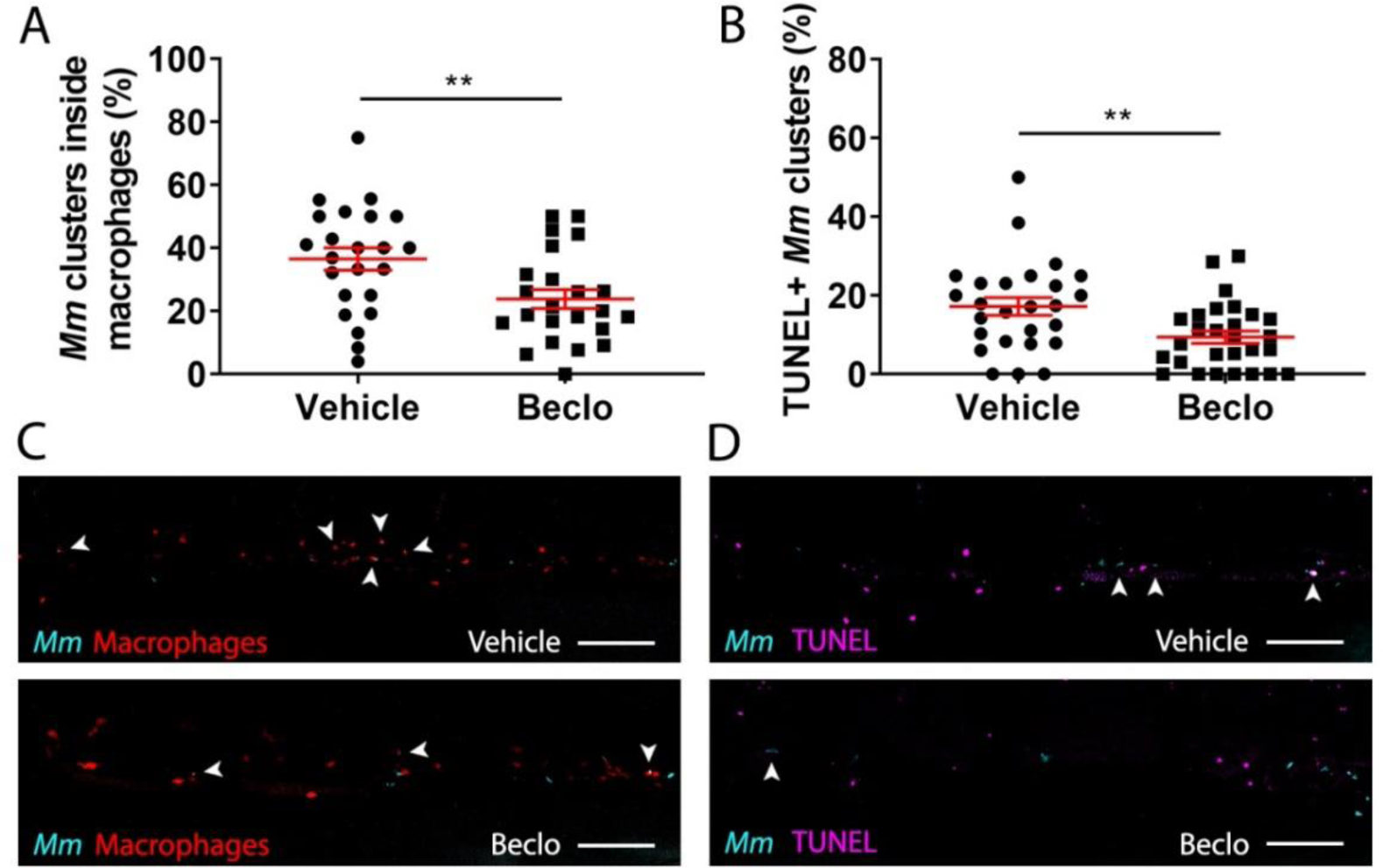
Effect of beclomethasone on intracellular bacterial growth and cell death. Infection was performed in *Tg(mpeg1:mCherry-F)* embryos at 28 hpf, a TUNEL assay was performed at 48 hpi, and the CHT region of the embryos was imaged using confocal microscopy. A. The percentage of *Mm* clusters that were inside macrophages based on colocalization with the red fluorescent signal from mCherry. Statistical analysis was performed by two-tailed t-test. In the beclomethasone-treated group, the percentage of *Mm* clusters inside macrophages was significantly lower compared to the vehicle-treated group. B. The percentage of TUNEL-positive *Mm* clusters. Statistical analysis by two-tailed t-test showed that the beclomethasone-treated group had a significantly lower percentage of TUNEL+ *Mm* clusters. In panels A and B, each data point represents a single embryo and the means ± s.e.m. of data accumulated from three independent experiments are shown in red. Statistical significance is indicated by: ** P<0.01. C. Representative confocal microscopy images of macrophage phagocytosis. Bacteria are shown in blue and macrophages in red. Arrowheads indicate intracellular bacterial clusters. Scale bar = 100 μm. D. Representative confocal microscopy images of cell death (TUNEL+ cells in magenta) and *Mm* infection (bacteria in blue). Arrowheads indicate bacterial clusters overlapping with TUNEL+ cells. Scale bar = 100 μm.

### Beclomethasone inhibits phagocytosis-related gene expression in macrophages

To unravel the molecular mechanisms underlying the beclomethasone-induced inhibition of the phagocytic activity of macrophages, we performed qPCR analysis on FACS-sorted macrophages derived from 28 hpf larvae after 2 h of beclomethasone treatment. To determine the phenotype of the sorted macrophages, the expression of a classic pro-inflammatory gene, *tnfa*, was measured (Martinez and Gordon, 2014; Nguyen-Chi et al., 2015). The levels of *tnfa* expression were significantly lower after beclomethasone treatment (Figure 6 A), in agreement with previously reported transcriptome analysis (Xie et al., 2019). In addition, we measured the expression levels of seven phagocytosis-related genes, *sparcl1, uchl1, ube2v1, marcksa, marcksb, bsg* and *tubb5* (Banerjee et al., 2019; Carballo et al., 1999; Jeon et al., 2010) (Figure 6 B-H). The expression levels of four of these genes, *sparcl1, uchl1, marcksa* and *marcksb*, were inhibited by beclomethasone treatment, while the levels of the other three (*ube2v1, bsg* and *tubb5)* were not affected. These data suggest that beclomethasone inhibits the phagocytic activity of macrophages by suppressing the transcription of phagocytosis-related genes in these cells.

**Figure 6.**
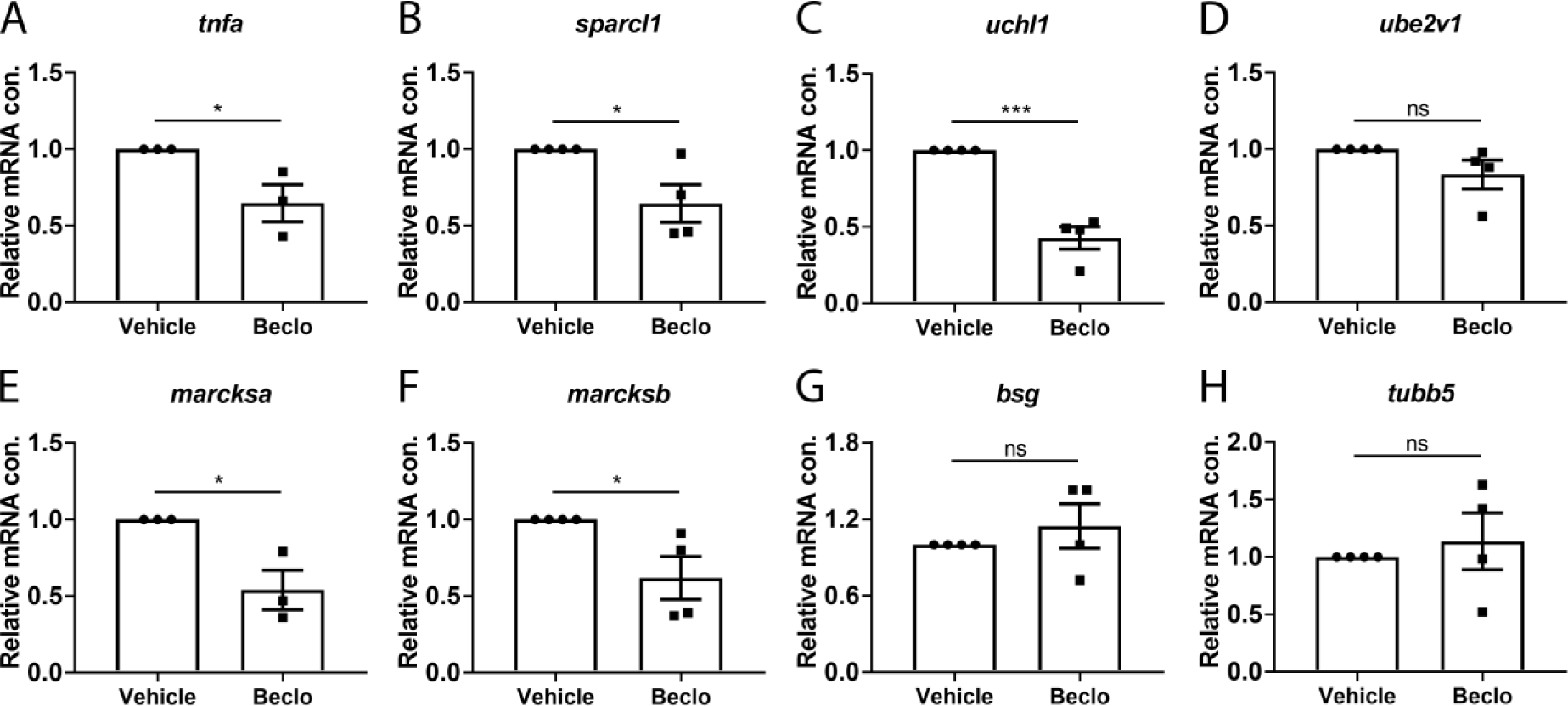
Effect of beclomethasone on gene expression levels in FACS-sorted macrophages. At 28 hpf, *Tg(mpeg1:mCherry-F)* embryos were treated with vehicle or beclomethasone for two hours, after which macrophages were isolated by FACS sorting. Gene expression levels were determined in the sorted cells by qPCR for *tnfa* (A), *sparcl1* (B), *uchl1* (C), *ube2v1* (D), *marcksa* (E), *marcksb* (F), *bsg* (G) and *tubb5* (H). Statistical analysis by two-tailed t-test showed that the levels of *tnfa, sparcl1, uchl1, marcksa* and *marcksb* expression were significantly inhibited by beclomethasone treatment. Data shown are the means ± s.e.m. of three or four independent experiments, and markers show averages of individual experiments. Statistical significance is indicated by: ns, non-significant; * P<0.05; *** P<0.001.

### Effect of Beclomethasone on the phagocytosis of *Salmonella* Typhimurium

To study whether the beclomethasone-induced inhibitory effect on macrophage phagocytosis of *Mm* can be generalized to other bacterial infections, we analyzed the effect of beclomethasone on infection with *Salmonella* Typhimurium, which is also an intracellular pathogen, but belongs to the gram-negative class. We quantified the percentages of bacteria phagocytosed by macrophages at different time points after infection in the *Tg(mpeg1:GFP)* fish line (Figure 7). In the vehicle group, the percentage of phagocytosed *Salmonella* Typhimurium increased from 5.7±0.7% at 10 mpi to 9.0±1.2% at 30 mpi and 17.9±1.7% at 60 mpi, and these percentages were significantly lower in the beclomethasone-treated group at all time points (3.1±0.5% at 10 mpi, 6.5±1.0% at 30 mpi and 10.0±1.4% at 60 mpi). These data demonstrate that the inhibitory effect of beclomethasone on the phagocytic activity of macrophages is not specific for *Mm*, but can also be observed for a distantly related *Salmonella* species.

**Figure 7.**
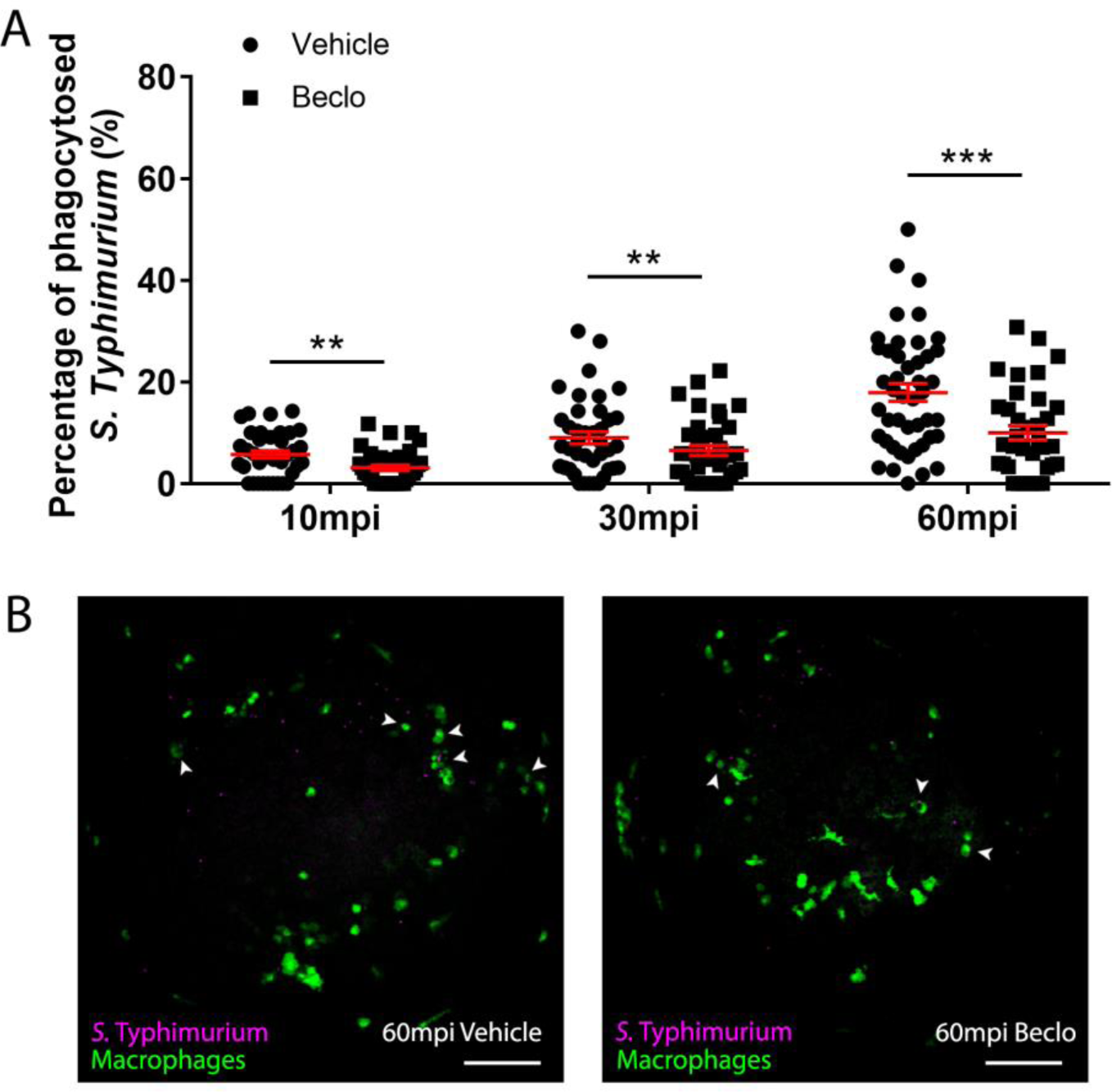
Effect of beclomethasone on phagocytosis of *Salmonella* Typhimurium. At 28 hpf *Tg(mpeg1:GFP)* embryos (vehicle- or beclomethasone-treated) were infected with *S*. Typhimurium through intravenous injection. At 10, 30, and 60 mpi, confocal microscopy images were taken of the yolk area, as indicated in Figure 4A, and the macrophage phagocytic capacity was determined. A. Percentage of phagocytosed *S*. Typhimurium at 10, 30 and 60 mpi. Statistical analysis, performed by fitting data to a beta inflated regression with Tukey’s post hoc test, showed that the phagocytic activity of macrophages was significantly inhibited by beclomethasone treatment at 60 mpi, and not at other time points. Each data point represents a single embryo and the means ± s.e.m. of data accumulated from three independent experiments are shown in red. Statistical significance is indicated by: ns, non-significant; **** P<0.0001. B. Representative confocal microscopy images of infected vehicle- and beclomethasone-treated individuals at 60 mpi. Bacteria are shown in magenta, macrophages in green. Arrowheads indicate bacteria phagocytosed by macrophages. Scale bar = 100 μm.

## Discussion

Synthetic GCs are widely prescribed to treat various immune-related diseases, but their clinical use is limited by the severe side effects evoked by prolonged therapy, including a higher susceptibility to TB (Caplan et al., 2017; Jick et al., 2006). In order to gain more insight into the mechanism underlying this GC effect, we used the zebrafish *Mm* infection model, which mimics human TB, and studied the effect of GC treatment on the development of the infection. We showed that GC treatment increased the level of *Mm* infection, which was reflected in the overall bacterial burden, the size and number of bacterial clusters and the level of dissemination. Since we found that GC treatment inhibited the phagocytic activity but not the microbicidal capacity of macrophages, we propose that the GC-induced increase in infection susceptibility is due to the inhibition on phagocytosis. Analysis of the transcription level of phagocytosis-related genes in macrophages suggested that the inhibition of phagocytic activity by GCs is mediated by Gr interfering with phagocytosis-related gene transcription. As a result of the lower phagocytic activity of the macrophages, the percentage of intracellular bacteria is decreased, which results in a lower level of cell death due to the *Mm* infection and exacerbated growth of the extracellular bacterial fraction. Finally, we showed that GC treatment not only limited phagocytosis of mycobacteria, but also of a Salmonella species, which suggests that the decrease in phagocytic activity may also explain the increased susceptibility to other bacterial infections that is commonly observed in patients receiving GC therapy (Caplan et al., 2017; Dixon et al., 2011; Fardet et al., 2016).

Upon bacterial infections, macrophages are the first responders of the immune system. In humans, *Mtb* generally infects lungs due to its air transmission properties and in the lungs it is taken up by alveolar macrophages within the first few days. In later stages, *Mtb* replicates, translocates to secondary loci and aggregates into granulomas with other attracted immune cells (Cambier et al., 2014; Russell, 2011; Srivastava et al., 2014). Consistently, in the zebrafish model, *Mm* is predominantly phagocytosed by macrophages within 30-60 min after intravenous infection in embryos, leading to initial stages of granuloma formation in the next few days (Benard et al., 2014; Davis et al., 2002). The phagocytosis activity and microbicidal capacity of macrophages have both been shown to be important for dealing with *Mm* infection (Benard et al., 2014; Clay et al., 2007). Interestingly, in our study we found that the microbicidal capacity of macrophages was not affected by GC treatment, which suggests that the inhibition of macrophage phagocytosis is a specific effect of GCs targeted at the uptake of pathogens rather than a global suppression of anti-microbial processes in macrophages.

Our study in the zebrafish model provides *in vivo* evidence for GC interference with macrophage phagocytosis that confirms results from various other studies. In line with our results, it has previously been shown that GCs decrease the phagocytosis of several *Escherichia coli* strains by human monocyte-derived (THP-1) macrophages and by murine bone marrow-derived macrophages (BMDMs) (Olivares-Morales et al., 2018). Similarly to our results, in this study the reduced phagocytosis activity was accompanied by a decreased expression of genes involved in phagosome formation including *MARCKS* and pro-inflammatory genes like *TNF* (Olivares-Morales et al., 2018). In earlier studies, decreased macrophage phagocytosis of carbon particles was observed *in vivo*, in GC-treated rats and rheumatoid arthritis patients (Jessop et al., 1973; Vernon-Roberts et al., 1973).

In contrast to our results on phagocytosis of mycobacteria, in other studies GC treatment has been shown to enhance the phagocytosis activity of macrophages. Upon GC exposure, increased phagocytosis of human monocyte-derived macrophages was observed for *Haemophillus influenzae* and *Streptococcus pneumoniae* (Taylor et al., 2010), and *Staphylococcus aureus* (van der Goes et al., 2000). This increased phagocytic activity would be in line with the well-established GC-induced enhancement of the phagocytosis of apoptotic neutrophils, which has been observed in, differentiated THP-1 macrophages, through stimulation of a protein S/Mer tyrosine kinase dependent pathway (Liu et al., 1999; McColl et al., 2009; Zahuczky et al., 2011), and in mouse alveolar macrophages (McCubbrey et al., 2012). This effect is considered to play an important role in GCs actively promoting the resolution of inflammation and reflects the GC-enhanced differentiation of macrophages to an anti-inflammatory phenotype (Busillo and Cidlowski, 2013; Ehrchen et al., 2019). Interestingly, GC treatment does not enhance the phagocytosis capacity in differentiated THP-1 macrophages of latex beads or apoptotic cells (Zahuczky et al., 2011). Most likely, the effects of GCs on the phagocytic activity of macrophages are highly dependent on the differentiation status of the cells, the particles they encounter and the tissue environment.

Our study revealed an inhibitory effect of GCs on four phagocytosis-related genes in FACS-sorted macrophages: *sparcl-1, uchl-1, marcksa and marcksb*. Among those genes, the human and mouse homologs of *sparcl-1* and *uchl-1* were reported to have a phagocytosis-promoting activity (Banerjee et al., 2019; Jeon et al., 2010). In human THP-1-derived macrophages, MARCKS plays a role in cytoskeletal remodeling and phagosome formation, and in line with our study the *MARCKS* gene expression was found to be inhibited by dexamethasone treatment (Carballo et al., 1999; Olivares-Morales et al., 2018). Together with our observation that phagocytosis is dependent on *de novo* protein synthesis, these results support the idea that GC treatment inhibits the phagocytosis activity of macrophages through interfering with transcription of genes that stimulate the phagocytic activity.

After internalization by macrophages, *Mm* are exposed to a bactericidal environment (Lesley and Ramakrishnan, 2008). Some bacteria may be killed by macrophages, while others may proliferate mediated by virulence determinants like Erp and RD1 (Clay et al., 2008; Lesley and Ramakrishnan, 2008; Lewis et al., 2003). When the macrophages are incapable of containing the bacteria, they undergo cell death leading to recruitment of more macrophages (Davis and Ramakrishnan, 2009). In our study, GC treatment led to a lower percentage of intracellular *Mm* at later stages, consistent with the decreased phagocytosis at early time points, and consequently less *Mm*-related cell death. The GC treatment may also directly affect cell death, since in a recent study it was demonstrated that GCs inhibit necrosis of various *Mtb* infected mouse and human cell types by activating MKP-1, which suppresses a pathway involving p38 MAPK activation ultimately leading to a loss of mitochondrial integrity (Gräb et al., 2019). The increased numbers of extracellular bacteria could traverse endothelial barriers directly and grow more rapidly in a less restrictive environment outside macrophages, which may explain our observation of a higher bacterial burden induced by GC treatment.

Based on our results, it may seem surprising that adjunctive GC therapy is often beneficial to TB patients, and even increases survival among tuberculous meningitis and pericarditis patients (Strang et al., 2004; Thwaites et al., 2004; Wiysonge et al., 2017). However, many of these observed beneficial effects are either minor or under debate. This may be due to GC therapy benefiting only a subset of patients whose disease has mainly progressed as a result of an excessive inflammatory response (which can be controlled with GC therapy), rather than a failed reaction to the infection, which was demonstrated for GC-treated TB meningitis patients with specific polymorphisms in the *LTA4H* gene (Tobin et al., 2012). We therefore suggest that in a subset of patients at later stages of infection, the anti-inflammatory effects of a GC treatment may outweigh a possible inhibitory effect on the phagocytic activity of the macrophages. Further research using the zebrafish model may shed light on a possible interplay between these effects, since the *Mm* infection model has been shown to have excellent translational value for human TB, including the effects of GC treatment (Tobin 2010, Tobin 2012).

In conclusion, our *in vivo* study on the effect of GC treatment in the zebrafish *Mm* infection model shows that GCs, through activation of Gr, inhibit the phagocytic activity of macrophages, which results in more extracellular bacterial growth and a higher infection level. These results may explain why clinically prolonged GC treatment is associated with an increased risk of TB and other bacterial infections.

## Materials and methods

### Zebrafish lines and maintenance

Zebrafish were maintained and handled according to the guidelines from the Zebrafish Model Organism Database (http://zfin.org) and in compliance with the directives of the local animal welfare body of Leiden University. They were exposed to a 14 hours light and 10 hours dark cycle to maintain circadian rhythmicity. Fertilization was performed by natural spawning at the beginning of the light period. Eggs were collected and raised at 28°C in egg water (60 µg/ml Instant Ocean sea salts and 0.0025% methylene blue). The following fish lines were used: wild type strain AB/TL, and the transgenic lines *Tg(mpeg1:mCherry*-*F*^*umsF001*^) (Bernut et al., 2014) and *Tg(mpeg1:eGFP*^*gl22*^) (Ellett et al., 2011).

### Bacterial culture and infection through intravenous injections

Bacteria used for this study were *Mycobacteria marinum*, strain M, constitutively fluorescently labelled with Wasabi or mCrimson (Ramakrishnan and Falkow, 1994; Takaki et al., 2013), *Mm* mutant *Δerp* labelled with Wasabi (Cosma et al., 2006), and *Salmonella enterica serovar* Typhimurium (*S*. Typhimurium) wild type (wt) strain SL1344 labelled with mCherry (Burton et al., 2014; Hoiseth and Stocker, 1981). The *Mm* strain M and *S*. Typhimurium wt strain were cultured at 28°C and 37°C respectively and the bacterial suspensions were prepared with phosphate buffered saline (PBS) with 2% (w/v) polyvinylpyrrolidone-40 (PVP40, Sigma-Aldrich), as previously described (Benard et al., 2012). The suspension of *Mm Δerp-*Wasabi was prepared directly from −80°C frozen aliquots.

After anesthesia with 0.02% aminobenzoic acid ethyl ester (tricaine, Sigma-Aldrich), 28 hours post fertilization (hpf) embryos were injected with *Mm* or *S*. Typhimurium into the blood island (or hindbrain if specified) under a Leica M165C stereomicroscope, as previously described (Benard et al., 2012). The injection dose was 200 CFU for *Mm* and 50 CFU for *S*. Typhimurium.

### Chemical treatments and bacterial burden quantification

The embryos were treated with 25 μM (or different if specified) beclomethasone (Sigma-Aldrich) or vehicle (0.05% dimethyl sulfoxide (DMSO)) in egg water from 2 hours before injection to the end of an experiment. RU-486 (Sigma-Aldrich) was administered at a concentration of 5 μM (0.02% DMSO), and cycloheximide (Sigma-Aldrich) at 100 μg/ml (0.04% DMSO). If the treatment lasted longer than 1 day, the medium was refreshed every day.

For bacterial burden quantification, the embryos from the vehicle- and beclomethasone-treated groups were imaged alive using a Leica M205FA fluorescence stereomicroscope equipped with a Leica DFC 345FX camera (Leica Microsystems). The images were analyzed using custom-designed pixel quantification software (previously described by Benard et al. (2015)), and Image J (plugin ‘Analyze Particles’).

### Hindbrain infection and analysis of dissemination

To assess the dissemination efficiency, the embryos were injected with 50 CFU *Mm* into the hindbrain at 28 hpf. At 2 dpi, the embryos were imaged with a Leica M205FA fluorescence stereomicroscope equipped with a Leica DFC 345FX camera. The embryos were classified into two categories: with or without disseminated infection. An embryo was considered without disseminated infection if all the bacteria were still contained in the hindbrain ventricle and considered with dissemination if bacteria were present in any other part of the embryo.

### Analysis of microbicidal activity

After infection at 28 hpf with *Mm Δerp-*Wasabi, *Tg(mpeg1:mCherry-F)* embryos were fixed at 44 hpi with 4% paraformaldehyde (PFA, Sigma-Aldrich) and imaged using a Leica TCS SP8 confocal microscope with 40X objective (NA 1.3). All macrophages that contained *Mm Δerp-*Wasabi in the tail region were analyzed. The level of infection inside macrophages was classified into two categories based on the number of bacteria: 1-10 bacteria or >10 bacteria, following established protocols (Clay et al., 2008; Sommer et al., 2020).

### Analysis of phagocytic activity

After infection at 28 hpf with *Mm*-Wasabi or *S*. Typhimurium-mCherry, *Tg(mpeg1:mCherry-F)* or *Tg(mpeg1:GFP)* embryos were fixed with 4% PFA at different time points and imaged using a Leica TCS SP8 confocal microscope with 20X objective (NA 0.75). The yolk sac area was selected as the quantification area (Figure 4A). The number of fluorescently labelled *Mm* or *S*. Typhimurium in this area, and those present inside a macrophage, were counted in a manual and blinded way.

### TUNEL assay

After infection at 28 hpf, *Tg(mpeg1:mCherry-F)* embryos were fixed with 4% PFA at 48 hpi and stained using terminal deoxynucleotidyl transferase dUTP nick end labelling (TUNEL) with the In Situ Cell Death Detection Kit, TMR red (Sigma-Aldrich), as previously described by Zhang et al. (2019). For this TUNEL staining, the embryos were first dehydrated and then rehydrated gradually with methanol in PBS, and permeabilized with 10 μg/ml Proteinase K (Roche). The embryos were subsequently fixed with 4% PFA for another 20 min and stained with reagent mixture overnight at 37°C. After the reaction was stopped by washing with PBS containing 0.05% Tween-20 (PBST), the CHT region of the embryos was imaged using a Leica TCS SP8 confocal microscope with 40X objective (NA 1.3). The total number of fluorescently labelled *Mm* clusters and the number of these clusters overlapping with TUNEL staining were counted in a manual and blinded way.

### Fluorescence-Activated Cell Sorting (FACS) of macrophages

Macrophages were sorted from *Tg(mpeg1:mCherry-F)* embryos as previously described (Rougeot et al., 2014; Zakrzewska et al., 2010). Dissociation was performed with 150-200 embryos for each sample after 2 hours beclomethasone or vehicle treatment (started at 28 hpf) using Liberase TL (Roche) and stopped by adding Fetal Calf Serum (FCS) to a final concentration of 10%. Isolated cells were resuspended in Dulbecco’s PBS (DPBS), and filtered through a 40 μm cell strainer. Actinomycin D (Sigma-Aldrich) was added (final concentration of 1 µg/ml) to each step to inhibit transcription. Macrophages were sorted based on their red fluorescent signal using a FACSAria III cell sorter (BD Biosciences). The sorted cells were collected in QIAzol lysis reagent (Qiagen) for RNA isolation.

### RNA isolation, cDNA synthesis and quantitative PCR (qPCR) analysis

RNA isolation from FACS-sorted cells was performed using the miRNeasy mini kit (Qiagen), according to the manufacturer’s instructions. Extracted total RNA was reverse-transcribed using the iScript(tm) cDNA Synthesis Kit (Bio-Rad). QPCR was performed on a MyiQ Single-Color Real-Time PCR Detection System (Bio-Rad) using iTaq(tm) Universal SYBR® Green Supermix (Bio-Rad). The sequences of the primers used are provided in Supplementary Table 1. Cycling conditions were pre-denaturation for 3 min at 95°C, followed by 40 cycles of denaturation for 15 s at 95°C, annealing for 30 s at 60°C, and elongation for 30 s at 72°C. Fluorescent signals were measured at the end of each cycle. Cycle threshold values (Ct values, i.e. the cycle numbers at which a threshold value of the fluorescence intensity was reached) were determined for each sample. To determine the gene regulation due to beclomethasone treatment in each experiment, the average Ct value of the beclomethasone treated samples was subtracted from the average Ct value of the vehicle-treated samples, and the fold change of gene expression was calculated, which was subsequently adjusted to the expression levels of a reference gene (*peptidylprolyl isomerase Ab* (*ppiab*)).

## Statistical analysis

Statistical analysis was performed using GraphPad Prism by one-way ANOVA with Bonferroni’s post hoc test (Figure 1A) or two-way ANOVA with Tukey’s post hoc test (Figure 1B, Figure 2 A-C) or two-tailed t-test (Figure 2D, Figure 3, 5, 6) or using R Statistical Software by fitting data to a beta inflated regression (from ‘gamlss’ package) (Stasinopoulos and Rigby, 2007) with Tukey’s post hoc test (Figure 4, 7).

## Acknowledgements

We thank Frida Sommer for her advice on *Mm Δerp* assays, Ralf Boland, Salomé Munoz Sanchez, Dr. Michiel van der Vaart and Aleksandra Fesliyska for their assistance with bacterial infections, Dr. Rui Zhang and Dr. Monica Varela Alvarez for their suggestions concerning TUNEL assays and Patrick van Hage for his help with the statistical analysis. We thank the fish facility team, in particular Ulrike Nehrdich, Ruth van Koppen, Karen Bosma and Guus van der Velden for zebrafish maintenance. We thank Dr. Georges Lutfalla and Dr. Graham Lieschke for providing transgenic zebrafish lines.

## References

Alsop, D. and Vijayan, M. M. (2008). Development of the corticosteroid stress axis and receptor expression in zebrafish. Am J Physiol Regul Integr Comp Physiol 294, R711–9.

Alzeer, A. H. and FitzGerald, J. M. (1993). Corticosteroids and tuberculosis: risks and use as adjunct therapy. Tuber Lung Dis 74, 6–11.

Banerjee, H., Krauss, C., Worthington, M., Banerjee, N., Walker, R. S., Hodges, S., Chen, L., Rawat, K., Dasgupta, S., Ghosh, S. et al. (2019). Differential expression of efferocytosis and phagocytosis associated genes in tumor associated macrophages exposed to African American patient derived prostate cancer microenvironment. J Solid Tumors 9, 22–27.

Benard, E. L., Roobol, S. J., Spaink, H. P. and Meijer, A. H. (2014). Phagocytosis of mycobacteria by zebrafish macrophages is dependent on the scavenger receptor Marco, a key control factor of pro-inflammatory signalling. Dev Comp Immunol 47, 223–33.

Benard, E. L., van der Sar, A. M., Ellett, F., Lieschke, G. J., Spaink, H. P. and Meijer, A. H. (2012). Infection of zebrafish embryos with intracellular bacterial pathogens. J Vis Exp.

Bernut, A., Herrmann, J.-L., Kissa, K., Dubremetz, J.-F., Gaillard, J.-L., Lutfalla, G. and Kremer, L. (2014). Mycobacterium abscessus cording prevents phagocytosis and promotes abscess formation. Proceedings of the National Academy of Sciences 111, E943–E952.

Bovornkitti, S., Kangsadal, P., Sathirapat, P. and Oonsombatti, P. (1960). Reversion and Reconversion Rate of Tuberculin Skin Reactions in Correlation with the Use of Prednisone<sup>1</sup>. Diseases of the Chest 38, 51–55.

Buckley, L. and Humphrey, M. B. (2018). Glucocorticoid-Induced Osteoporosis. N Engl J Med 379, 2547–2556.

Burton, N. A., Schürmann, N., Casse, O., Steeb, A. K., Claudi, B., Zankl, J., Schmidt, A. and Bumann, D. (2014). Disparate impact of oxidative host defenses determines the fate of Salmonella during systemic infection in mice. Cell Host & Microbe 15, 72–83.

Busillo, J. M. and Cidlowski, J. A. (2013). The five Rs of glucocorticoid action during inflammation: ready, reinforce, repress, resolve, and restore. Trends Endocrinol Metab 24, 109–19.

Cambier, C. J., Falkow, S. and Ramakrishnan, L. (2014). Host evasion and exploitation schemes of Mycobacterium tuberculosis. Cell 159, 1497–509.

Caplan, A., Fett, N., Rosenbach, M., Werth, V. P. and Micheletti, R. G. (2017). Prevention and management of glucocorticoid-induced side effects: A comprehensive review: A review of glucocorticoid pharmacology and bone health. J Am Acad Dermatol 76, 1–9.

Carballo, E., Pitterle, D. M., Stumpo, D. J., Sperling, R. T. and Blackshear, P. J. (1999). Phagocytic and macropinocytic activity in MARCKS-deficient macrophages and fibroblasts. American Journal of Physiology-Cell Physiology 277, C163–C173.

Chatzopoulou, A., Heijmans, J. P., Burgerhout, E., Oskam, N., Spaink, H. P., Meijer, A. H. and Schaaf, M. J. (2016). Glucocorticoid-induced attenuation of the inflammatory response in zebrafish. Endocrinology 157, 2772–2784.

Chatzopoulou, A., Roy, U., Meijer, A. H., Alia, A., Spaink, H. P. and Schaaf, M. J. (2015). Transcriptional and metabolic effects of glucocorticoid receptor α and β signaling in zebrafish. Endocrinology 156, 1757–1769.

Clay, H., Davis, J. M., Beery, D., Huttenlocher, A., Lyons, S. E. and Ramakrishnan, L. (2007). Dichotomous role of the macrophage in early Mycobacterium marinum infection of the zebrafish. Cell Host Microbe 2, 29–39.

Clay, H., Volkman, H. E. and Ramakrishnan, L. (2008). Tumor necrosis factor signaling mediates resistance to mycobacteria by inhibiting bacterial growth and macrophage death. Immunity 29, 283–94.

Cosma, C. L., Klein, K., Kim, R., Beery, D. and Ramakrishnan, L. (2006). Mycobacterium marinum Erp is a virulence determinant required for cell wall integrity and intracellular survival. Infection and immunity 74, 3125–3133.

Critchley, J. A., Orton, L. C. and Pearson, F. (2014). Adjunctive steroid therapy for managing pulmonary tuberculosis. Cochrane Database Syst Rev, Cd011370.

Cronan, M. R. and Tobin, D. M. (2014). Fit for consumption: zebrafish as a model for tuberculosis. Disease models & mechanisms 7, 777–784.

Davis, J. M., Clay, H., Lewis, J. L., Ghori, N., Herbomel, P. and Ramakrishnan, L. (2002). Real-Time Visualization of Mycobacterium-Macrophage Interactions Leading to Initiation of Granuloma Formation in Zebrafish Embryos. Immunity 17, 693–702.

Davis, J. M. and Ramakrishnan, L. (2009). The Role of the Granuloma in Expansion and Dissemination of Early Tuberculous Infection. Cell 136, 37–49.

Dixon, W., Kezouh, A., Bernatsky, S. and Suissa, S. (2011). The influence of systemic glucocorticoid therapy upon the risk of non-serious infection in older patients with rheumatoid arthritis: a nested case–control study. Annals of the rheumatic diseases 70, 956–960.

Drain, P. K., Bajema, K. L., Dowdy, D., Dheda, K., Naidoo, K., Schumacher, S. G., Ma, S., Meermeier, E., Lewinsohn, D. M. and Sherman, D. R. (2018). Incipient and Subclinical Tuberculosis: a Clinical Review of Early Stages and Progression of Infection. Clin Microbiol Rev 31.

Ehrchen, J. M., Roth, J. and Barczyk-Kahlert, K. (2019). More Than Suppression: Glucocorticoid Action on Monocytes and Macrophages. Front Immunol 10, 2028.

Ellett, F., Pase, L., Hayman, J. W., Andrianopoulos, A. and Lieschke, G. J. (2011). mpeg1 promoter transgenes direct macrophage-lineage expression in zebrafish. Blood 117, e49–e56.

Evans, D. J. (2008). The use of adjunctive corticosteroids in the treatment of pericardial, pleural and meningeal tuberculosis: Do they improve outcome? Respiratory Medicine 102, 793–800.

Facchinello, N., Skobo, T., Meneghetti, G., Colletti, E., Dinarello, A., Tiso, N., Costa, R., Gioacchini, G., Carnevali, O. and Argenton, F. (2017). nr3c1 null mutant zebrafish are viable and reveal DNA-binding-independent activities of the glucocorticoid receptor. Scientific reports 7, 1–13.

Fardet, L., Petersen, I. and Nazareth, I. (2016). Common Infections in Patients Prescribed Systemic Glucocorticoids in Primary Care: A Population-Based Cohort Study. PLoS Med 13, e1002024.

Faught, E. and Vijayan, M. M. (2019). Loss of the glucocorticoid receptor in zebrafish improves muscle glucose availability and increases growth. Am J Physiol Endocrinol Metab 316, E1093–e1104.

Furin, J., Cox, H. and Pai, M. (2019). Tuberculosis. The Lancet 393, 1642–1656.

Gräb, J., Suárez, I., van Gumpel, E., Winter, S., Schreiber, F., Esser, A., Hölscher, C., Fritsch, M., Herb, M. and Schramm, M. (2019). Corticosteroids inhibit Mycobacterium tuberculosis-induced necrotic host cell death by abrogating mitochondrial membrane permeability transition. Nature communications 10, 1–14.

Hawn, T. R., Matheson, A. I., Maley, S. N. and Vandal, O. (2013). Host-directed therapeutics for tuberculosis: can we harness the host? Microbiol. Mol. Biol. Rev. 77, 608–627.

Hoiseth, S. K. and Stocker, B. (1981). Aromatic-dependent Salmonella typhimurium are non-virulent and effective as live vaccines. Nature 291, 238–239.

Houben, R. M. and Dodd, P. J. (2016). The global burden of latent tuberculosis infection: a re-estimation using mathematical modelling. PLoS Med 13, e1002152.

Jeon, H., Go, Y., Seo, M., Lee, W. H. and Suk, K. (2010). Functional selection of phagocytosis-promoting genes: cell sorting-based selection. J Biomol Screen 15, 949–55.

Jessop, J., Vernon-Roberts, B. and Harris, J. (1973). Effects of gold salts and prednisolone on inflammatory cells. I. Phagocytic activity of macrophages and polymorphs in inflammatory exudates studied by a” skin-window” technique in rheumatoid and control patients. Annals of the rheumatic diseases 32, 294.

Jick, S. S., Lieberman, E. S., Rahman, M. U. and Choi, H. K. (2006). Glucocorticoid use, other associated factors, and the risk of tuberculosis. Arthritis Rheum 55, 19–26.

Kadhiravan, T. and Deepanjali, S. (2010). Role of corticosteroids in the treatment of tuberculosis: an evidence-based update. Indian J Chest Dis Allied Sci 52, 153–8.

Kim, H., Yoo, C., Baek, H., Lee, E. B., Ahn, C., Han, J., Kim, S., Lee, J.-S., Choe, K. and Song, Y.-W. (1998). Mycobacterium tuberculosis infection in a corticosteroid-treated rheumatic disease patient population. Clinical and experimental rheumatology 16, 9–13.

Kulchavenya, E. (2014). Extrapulmonary tuberculosis: are statistical reports accurate? Therapeutic Advances in Infectious Disease 2, 61–70.

Lerner, B. H. (1996). Can stress cause disease? Revisiting the tuberculosis research of Thomas Holmes, 1949-1961. Ann Intern Med 124, 673–80.

Lesley, R. and Ramakrishnan, L. (2008). Insights into early mycobacterial pathogenesis from the zebrafish. Curr Opin Microbiol 11, 277–83.

Lewis, K. N., Liao, R., Guinn, K. M., Hickey, M. J., Smith, S., Behr, M. A. and Sherman, D. R. (2003). Deletion of RD1 from Mycobacterium tuberculosis mimics bacille Calmette-Guerin attenuation. J Infect Dis 187, 117–23.

Lin, P. L. and Flynn, J. L. (2010). Understanding latent tuberculosis: a moving target. The Journal of Immunology 185, 15–22.

Liu, Y., Cousin, J. M., Hughes, J., Van Damme, J., Seckl, J. R., Haslett, C., Dransfield, I., Savill, J. and Rossi, A. G. (1999). Glucocorticoids Promote Nonphlogistic Phagocytosis of Apoptotic Leukocytes. The Journal of Immunology 162, 3639–3646.

Martinez, F. O. and Gordon, S. (2014). The M1 and M2 paradigm of macrophage activation: time for reassessment. F1000Prime Rep 6, 13.

McColl, A., Bournazos, S., Franz, S., Perretti, M., Morgan, B. P., Haslett, C. and Dransfield, I. (2009). Glucocorticoids induce protein S-dependent phagocytosis of apoptotic neutrophils by human macrophages. J Immunol 183, 2167–75.

McCubbrey, A. L., Sonstein, J., Ames, T. M., Freeman, C. M. and Curtis, J. L. (2012). Glucocorticoids relieve collectin-driven suppression of apoptotic cell uptake in murine alveolar macrophages through downregulation of SIRPα. The Journal of Immunology 189, 112–119.

Meijer, A. H. (2016). Protection and pathology in TB: learning from the zebrafish model. Semin Immunopathol 38, 261–73.

Muthuswamy, P., Hu, T.-C., Carasso, B., Antonio, M. and Dandamudi, N. (1995). Prednisone as adjunctive therapy in the management of pulmonary tuberculosis: report of 12 cases and review of the literature. CHEST 107, 1621–1630.

Nguyen-Chi, M., Laplace-Builhe, B., Travnickova, J., Luz-Crawford, P., Tejedor, G., Phan, Q. T., Duroux-Richard, I., Levraud, J. P., Kissa, K., Lutfalla, G. et al. (2015). Identification of polarized macrophage subsets in zebrafish. Elife 4, e07288.

Olivares-Morales, M. J., De La Fuente, M. K., Dubois-Camacho, K., Parada, D., Diaz-Jiménez, D., Torres-Riquelme, A., Xu, X., Chamorro-Veloso, N., Naves, R., Gonzalez, M.-J. et al. (2018). Glucocorticoids Impair Phagocytosis and Inflammatory Response Against Crohn’s Disease-Associated Adherent-Invasive Escherichia coli. Frontiers in Immunology 9.

Parikka, M., Hammaren, M. M., Harjula, S. K., Halfpenny, N. J., Oksanen, K. E., Lahtinen, M. J., Pajula, E. T., Iivanainen, A., Pesu, M. and Ramet, M. (2012). Mycobacterium marinum causes a latent infection that can be reactivated by gamma irradiation in adult zebrafish. PLoS Pathog 8, e1002944.

Prasad, K., Singh, M. B. and Ryan, H. (2016). Corticosteroids for managing tuberculous meningitis. Cochrane Database of Systematic Reviews.

Ramakrishnan, L. (2013). The zebrafish guide to tuberculosis immunity and treatment. Cold Spring Harb Symp Quant Biol 78, 179–92.

Ramakrishnan, L. and Falkow, S. (1994). Mycobacterium marinum persists in cultured mammalian cells in a temperature-restricted fashion. Infection and immunity 62, 3222–3229.

Rougeot, J., Zakrzewska, A., Kanwal, Z., Jansen, H. J., Spaink, H. P. and Meijer, A. H. (2014). RNA sequencing of FACS-sorted immune cell populations from zebrafish infection models to identify cell specific responses to intracellular pathogens. Methods Mol Biol 1197, 261–74.

Russell, D. G. (2011). Mycobacterium tuberculosis and the intimate discourse of a chronic infection. Immunological reviews 240, 252–268.

Ryan, H., Yoo, J. and Darsini, P. (2017). Corticosteroids for tuberculous pleurisy. Cochrane Database of Systematic Reviews.

Schaaf, M., Chatzopoulou, A. and Spaink, H. (2009). The zebrafish as a model system for glucocorticoid receptor research. Comparative Biochemistry and Physiology Part A: Molecular & Integrative Physiology 153, 75–82.

Schatz, M., Patterson, R., Kloner, R. and Falk, J. (1976). The prevalence of tuberculosis and positive tuberculin skin tests in a steroid-treated asthmatic population. Ann Intern Med 84, 261–5.

Singh, S. and Tiwari, K. (2017). Use of corticosteroids in tuberculosis. The Journal of Association of Chest Physicians 5, 70–75.

Smego, R. A. and Ahmed, N. (2003). A systematic review of the adjunctive use of systemic corticosteroids for pulmonary tuberculosis. Int J Tuberc Lung Dis 7, 208–13.

Sommer, F., Torraca, V., Kamel, S. M., Lombardi, A. and Meijer, A. H. (2020). Frontline Science: Antagonism between regular and atypical Cxcr3 receptors regulates macrophage migration during infection and injury in zebrafish. Journal of leukocyte biology 107, 185–203.

Srivastava, S., Ernst, J. D. and Desvignes, L. (2014). Beyond macrophages: the diversity of mononuclear cells in tuberculosis. Immunol Rev 262, 179–92.

Stasinopoulos, D. M. and Rigby, R. A. (2007). Generalized additive models for location scale and shape (GAMLSS) in R. Journal of Statistical Software 23, 1–46.

Stolte, E. H., van Kemenade, B. L. V., Savelkoul, H. F. and Flik, G. (2006). Evolution of glucocorticoid receptors with different glucocorticoid sensitivity. Journal of Endocrinology 190, 17–28.

Strang, J. I., Nunn, A. J., Johnson, D. A., Casbard, A., Gibson, D. G. and Girling, D. J. (2004). Management of tuberculous constrictive pericarditis and tuberculous pericardial effusion in Transkei: results at 10 years follow-up. Qjm 97, 525–35.

Suh, S. and Park, M. K. (2017). Glucocorticoid-Induced Diabetes Mellitus: An Important but Overlooked Problem. Endocrinol Metab (Seoul) 32, 180–189.

Takaki, K., Davis, J. M., Winglee, K. and Ramakrishnan, L. (2013). Evaluation of the pathogenesis and treatment of Mycobacterium marinum infection in zebrafish. Nature protocols 8, 1114.

Taylor, A., Finney-Hayward, T., Quint, J., Thomas, C., Tudhope, S., Wedzicha, J., Barnes, P. and Donnelly, L. (2010). Defective macrophage phagocytosis of bacteria in COPD. European Respiratory Journal 35, 1039–1047.

Thwaites, G. E., Nguyen, D. B., Nguyen, H. D., Hoang, T. Q., Do, T. T., Nguyen, T. C., Nguyen, Q. H., Nguyen, T. T., Nguyen, N. H., Nguyen, T. N. et al. (2004). Dexamethasone for the treatment of tuberculous meningitis in adolescents and adults. N Engl J Med 351, 1741–51.

Tobin, D. M. and Ramakrishnan, L. (2008). Comparative pathogenesis of Mycobacterium marinum and Mycobacterium tuberculosis. Cellular microbiology 10, 1027–1039.

Tobin, D. M., Roca, F. J., Oh, S. F., McFarland, R., Vickery, T. W., Ray, J. P., Ko, D. C., Zou, Y., Bang, N. D. and Chau, T. T. (2012). Host genotype-specific therapies can optimize the inflammatory response to mycobacterial infections. Cell 148, 434–446.

Tobin, D. M., Vary Jr, J. C., Ray, J. P., Walsh, G. S., Dunstan, S. J., Bang, N. D., Hagge, D. A., Khadge, S., King, M.-C. and Hawn, T. R. (2010). The lta4h locus modulates susceptibility to mycobacterial infection in zebrafish and humans. Cell 140, 717–730.

Torok, M. E., Nguyen, D. B., Tran, T. H., Nguyen, T. B., Thwaites, G. E., Hoang, T. Q., Nguyen, H. D., Tran, T. H., Nguyen, T. C., Hoang, H. T. et al. (2011). Dexamethasone and long-term outcome of tuberculous meningitis in Vietnamese adults and adolescents. PLoS One 6, e27821.

van der Goes, A., Hoekstra, K., van den Berg, T. K. and Dijkstra, C. D. (2000). Dexamethasone promotes phagocytosis and bacterial killing by human monocytes/macrophages in vitro. Journal of leukocyte biology 67, 801–807.

van Leeuwen, L. M., van der Sar, A. M. and Bitter, W. (2015). Animal models of tuberculosis: zebrafish. Cold Spring Harbor perspectives in medicine 5, a018580.

Vernon-Roberts, B., Jessop, J. and Dore, J. (1973). Effects of gold salts and prednisolone on inflammatory cells. II. Suppression of inflammation and phagocytosis in the rat. Annals of the rheumatic diseases 32, 301.

Wiysonge, C. S., Ntsekhe, M., Thabane, L., Volmink, J., Majombozi, D., Gumedze, F., Pandie, S. and Mayosi, B. M. (2017). Interventions for treating tuberculous pericarditis. Cochrane Database Syst Rev 9, Cd000526.

World Health Organization. (2019). Global tuberculosis report 2019. Geneva: World Health Organization.

Xie, Y., Tolmeijer, S., Oskam, J. M., Tonkens, T., Meijer, A. H. and Schaaf, M. J. (2019). Glucocorticoids inhibit macrophage differentiation towards a pro-inflammatory phenotype upon wounding without affecting their migration. Disease models & mechanisms 12, dmm037887.

Zahuczky, G., Kristóf, E., Majai, G. and Fésüs, L. (2011). Differentiation and Glucocorticoid Regulated Apopto-Phagocytic Gene Expression Patterns in Human Macrophages. Role of Mertk in Enhanced Phagocytosis. PLoS One 6, e21349.

Zakrzewska, A., Cui, C., Stockhammer, O. W., Benard, E. L., Spaink, H. P. and Meijer, A. H. (2010). Macrophage-specific gene functions in Spi1-directed innate immunity. Blood 116, e1–11.

Zhang, R., Varela, M., Forn-Cuni, G., Torraca, V., van der Vaart, M. and Meijer, A. H. (2019). Deficiency in the autophagy modulator Dram1 exacerbates pyroptotic cell death of Mycobacteria-infected macrophages. bioRxiv, 599266.

